# From Bile Acids to a Gas-Producing Microbiome Phenotype: A Novel Mechanism of Host–Microbiome Communication

**DOI:** 10.64898/2026.08.24.746699

**Authors:** M. Strus, T. Kasperski, K. Mech, A. Szczepanik, E. Golińska

## Abstract

**Background:** Microbiome-derived metabolites regulate host physiology, yet bacterial gaseous metabolites remain largely overlooked. Traditionally regarded as fermentation end-products, bacterial gases may act as biologically active mediators of host–microbiome communication. We hypothesized that bile acids regulate bacterial gaseous metabolism and influence host epithelial responses.

**Methods:** A high gas-producing clinical *Escherichia coli* isolate from a patient with moderately severe acute pancreatitis was cultured with selected primary and secondary bile acids. Gas production was assessed by pressure measurements, GC–TCD and GC–MS. Biological activity was evaluated by indirect exposure of Caco-2 and PANC-1 epithelial cells, followed by apoptosis/necrosis assays and whole-transcriptome RNA sequencing.

**Results:** Bile acids markedly reshaped bacterial gaseous metabolism. Cholic acid and deoxycholic acid promoted intense gas production, whereas chenodeoxycholic acid almost completely abolished it. Despite minimal apoptosis and necrosis, bacterial gaseous metabolites induced extensive transcriptional remodeling. Caco-2 cells showed stronger responses than PANC-1 cells, particularly to deoxycholic acid-derived gases, involving inflammatory signaling, extracellular matrix remodeling, epithelial plasticity, stress responses, and cancer-associated genes including **PTGS2, MMP1, PLAUR, NR4A2**, and **SERPINE1**. PANC-1 cells exhibited a more restricted response involving oxidative stress, proteostasis, and autophagy-associated pathways.

**Conclusions:** Our findings indicate that bacterial gases are a previously underrecognized class of microbiome-derived signaling molecules capable of modulating host gene expression independently of direct bacterial contact. We identify a **gas-producing microbiome phenotype** regulated by bile acid composition, linking microbial metabolism with epithelial signaling. These findings expand the concept of host–microbiome communication and provide a framework for investigating bacterial gaseous metabolites in intestinal and pancreatic diseases.

**Importance:** The gut microbiome produces numerous substances that can influence human health, but most research has focused on soluble metabolites such as short-chain fatty acids and bile acid derivatives. Bacterial gases, including hydrogen and carbon dioxide, have largely been considered metabolic waste products. Our study shows that this view may be incomplete. We demonstrate that bile acids can substantially change the amount and composition of gases produced by *Escherichia coli*, and that these bacterial gases can alter gene activity in human intestinal and pancreatic cells without direct bacterial contact. These findings identify bacterial gaseous metabolites as a previously underrecognized component of microbiome– host communication and suggest that differences in bacterial gas production may contribute to variation in epithelial responses along the gastrointestinal tract.

## 1. Introduction

The human gut microbiota is increasingly recognized as a key factor strongly linked to numerous diseases through the production of active compounds such as short-chain fatty acids (SCFAs), enzymes, lipopolysaccharides (LPS), vitamins, and numerous other bioactive metabolites, which influence epithelial barrier integrity, immune homeostasis, energy metabolism, and the function of many organs, including the pancreas (Fogelson et al. 2023; Pan et al. 2024). Although significant progress has been made in understanding the biological functions of these microbial products, bacterial gaseous metabolites remain among the least studied components of microbiota-derived signaling.

Hydrogen, carbon dioxide, methane, hydrogen sulfide, and other volatile compounds are constantly produced during bacterial carbohydrate fermentation and amino acid metabolism. Historically regarded as inert metabolic by-products of metabolism, these gases are now increasingly recognized as biologically active molecules capable of modulating oxidative stress, inflammatory responses, cellular metabolism, and intercellular communication (Pan et al. 2024; Wang et al. 2024). Several gaseous metabolites, particularly hydrogen, carbon dioxide, and hydrogen sulfide, readily diffuse across biological membranes, suggesting they may affect not only the intestinal epithelium but also distant organs. However, despite their potential physiological importance, the contribution of bacterial gas metabolites to pancreatic diseases remains largely unexplored.

Acute pancreatitis (AP) is characterized by profound disruption of intestinal barrier function, dysbiosis, bacterial translocation, and disturbances in bile acid metabolism. Accumulating clinical and experimental evidence indicates that the gut microbiota and microbial metabolites contribute to the severity and progression of the disease (Jia et al. 2024; Pan et al. 2024). Interestingly, imaging studies have occasionally demonstrated gas accumulation within the pancreas, peripancreatic tissues or biliary tract in patients with severe pancreatitis, even in the absence of gastrointestinal perforation or emphysematous infection.

The origin, chemical composition and biological significance of these gaseous accumulations remain largely unknown. In particular, it is unclear whether they originate exclusively from bacterial metabolism or whether they actively contribute to pancreatic inflammation. Bile acids may be a key factor regulating this process. In addition to their well-established role in lipid digestion, bile acids act as signaling molecules that shape both host metabolism and microbial physiology.

Moreover, intestinal bacteria actively modify bile acid composition through bile salt hydrolase activity and subsequent biotransformation reactions, thereby establishing a bidirectional interaction between bile acids and the gut microbiota.

Primary and secondary bile acids differ significantly in their antimicrobial activity and their ability to modulate bacterial growth, adaptation to stress, and metabolic pathways. *Enterobacterales*, especially *Escherichia coli*, exhibit remarkable metabolic plasticity, allowing adaptation to changing bile acid composition. However, the influence of individual primary and secondary bile acids on bacterial gas metabolism has not been systematically studied (Fogelson et al. 2023).

In our previous study, we isolated three strains of *Escherichia coli* from rectal swabs, stoma contents, and stool samples collected from a patient with moderately severe acute pancreatitis (MSAP). The isolates exhibited significant genotypic and phenotypic heterogeneity, particularly in their ability to produce extracellular gaseous metabolites in the presence of bile salts. One isolate, designated *E.coli* 33, exhibited a distinct gas-producing phenotype (Chmielarczyk et al. 2025). These findings suggest that bacterial gas metabolism is highly strain-dependent and strongly influenced by the presence of bile salts in the growth medium. Because gut bacteria rapidly deconjugate bile salts through bile salt hydrolase activity, generating free primary and secondary bile acids, we hypothesized that individual bile acids could regulate the qualitative and quantitative composition of bacterial gaseous metabolites throughout the gastrointestinal tract, thereby modifying microbiota-host interactions.

To test this hypothesis, we investigated the effects of primary and secondary bile acids on the quantity, composition, and pressure of gaseous metabolites produced by *E. coli* strain 33. Furthermore, we examined whether exposure to these bacterial gaseous metabolites modulates apoptosis, necrosis and global gene expression in human intestinal epithelial (Caco-2) and pancreatic epithelial (PANC-1) cells.

We further hypothesized that bile acid-dependent alterations in bacterial gaseous metabolism may represent a previously unrecognized mechanism of microbiota–host communication, linking intestinal dysbiosis with epithelial responses in both the gut and pancreas.

## 2. Materials and Methods

### 2.1. Bacterial isolates

The *Escherichia coli* strain EC33 used in the present study was selected based on the results of our previous clinical investigation, in which a collection of bacterial isolates was obtained from a single patient with moderately severe acute pancreatitis (MSAP). Among these isolates, three phenotypically distinct *E. coli* strains (EC18, EC25, and EC33) were identified and characterized. Briefly, MSAP developed during the patient’s hospitalization at the University Hospital of the Jagiellonian University Medical College, and microbiological specimens were collected as part of routine diagnostic procedures following approval by the Bioethics Committee of the Jagiellonian University (KBET1072.6120.279). Detailed clinical characteristics of the patient, microbiological findings, and the isolation procedure have been reported previously (Chmielarczyk et al., 2025). In our previous study, the three *E. coli* isolates exhibited marked phenotypic heterogeneity in extracellular gaseous metabolite production under bile-containing culture conditions. Notably, only strain EC33 consistently demonstrated a unique high gas-producing phenotype, clearly distinguishing it from the remaining isolates. Owing to this exceptional phenotype, EC33 was selected as the experimental model for all subsequent analyses performed in the present study.

### 2.2. Bacterial culture in the presence of primary and secondary bile acids

To investigate the influence of individual primary and secondary bile acids on the gaseous metabolism of the high gas-producing EC33 strain, the experimental procedure was based on our previously described bacterial gas production model (Chmielarczyk et al., 2025), with minor modifications.

Briefly, a 24-h culture of *E. coli* EC33 (approximately 1 × 10^8^ CFU/mL) was inoculated onto 1.75 L of solid MacConkey medium (Oxoid Ltd., Basingstoke, UK) prepared in sterile 2-L glass flasks.

The culture medium was supplemented separately with individual primary or secondary bile acids (Table 1) to evaluate their specific effects on the quantity and composition of gaseous metabolites produced by the high gas-producing EC33 strain. Control cultures were prepared under identical conditions without bile acid supplementation.

**Table 1.**
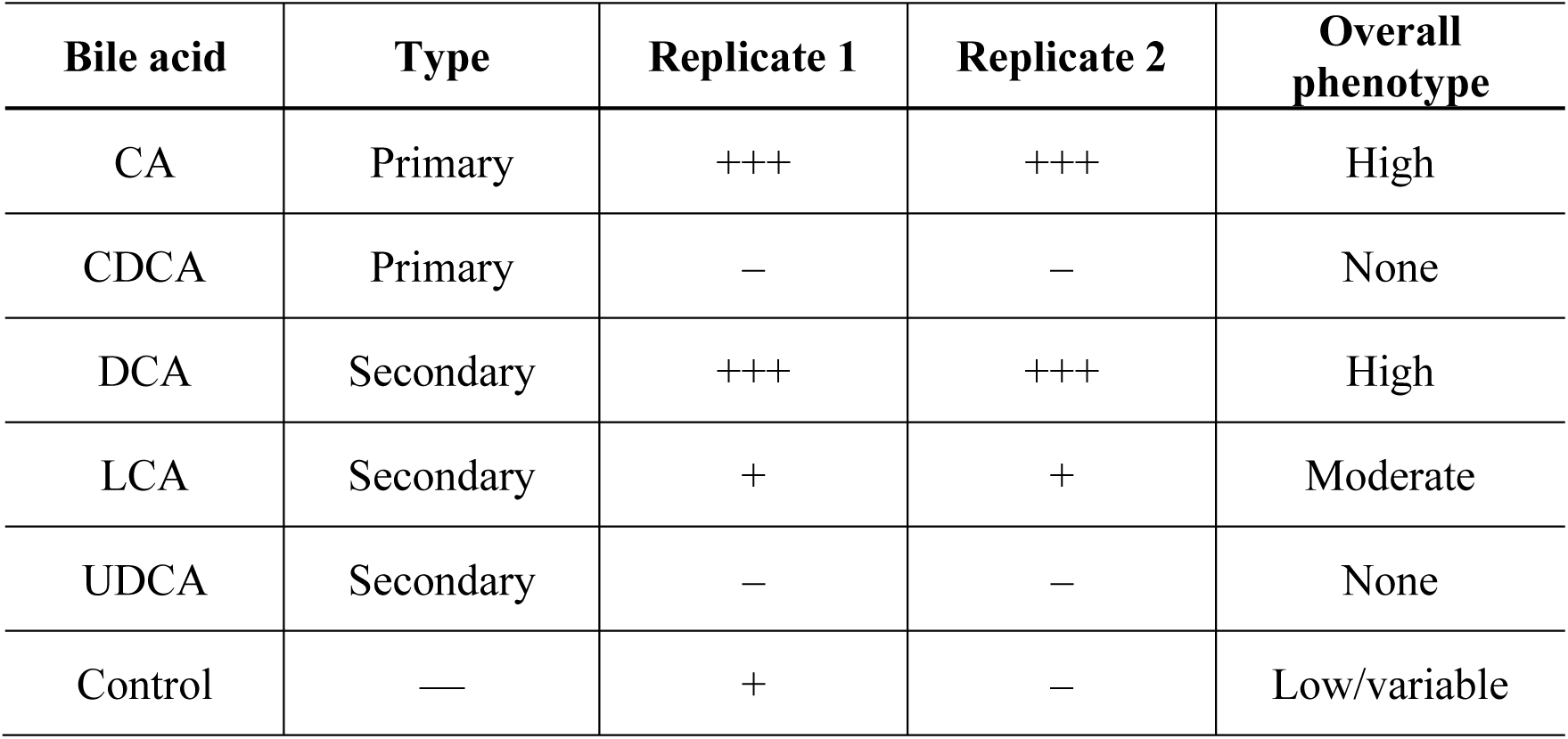
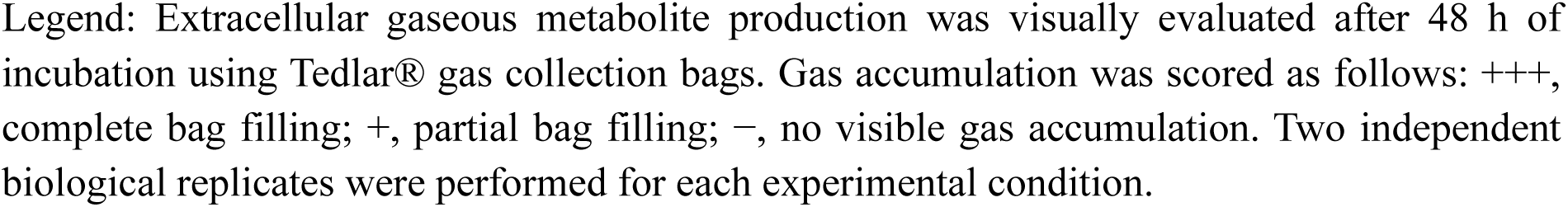
Visual assessment of the gas-producing phenotype of EC33 cultured in the presence of individual bile acids.

The flasks were connected to sterile 1-L Tedlar® gas sampling bags (Jensen Inert Products, Coral Springs, FL, USA), allowing continuous collection of bacterial gaseous metabolites released during incubation (Figure 1). All cultures were incubated aerobically at 37°C for 48 h. Based on our previous observations, differences in bacterial gas production between individual bile acid conditions became evident after 48 h of incubation; therefore, this time point was selected for all subsequent analyses. Each experimental condition was performed in duplicate.

**Figure 1.**
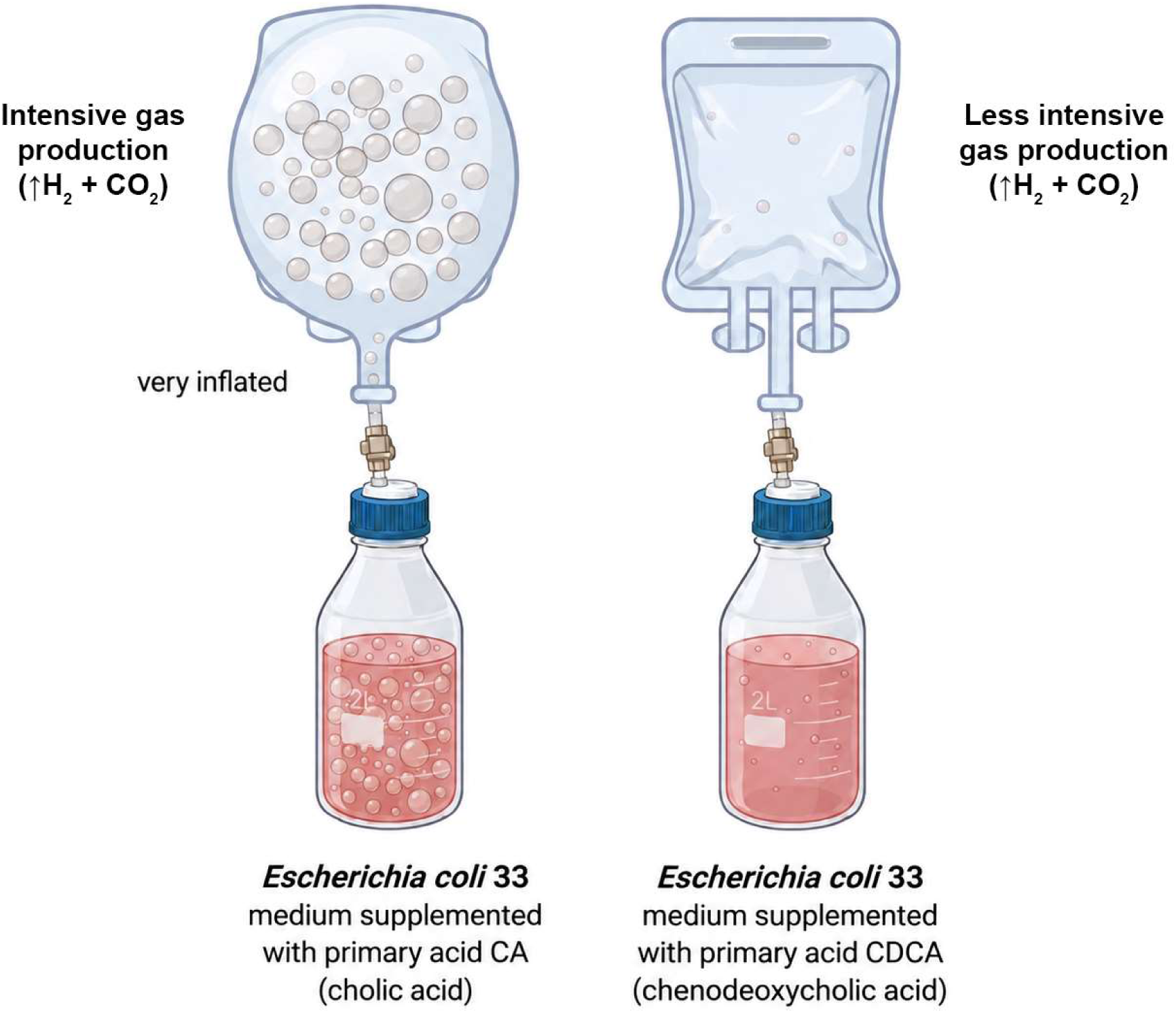
Schematic representation of the experimental setup used for bacterial gas collection. Bacterial cultures were grown in sealed 2L flasks connected via valves to Tedlar gas sampling bags. A 600 mL headspace was purged with air.

### 2.3. Quantitative and qualitative analysis of gaseous bacterial metabolites induced by primary and secondary bile acids

The composition and quantity of gaseous metabolites were determined using a GC–MS QP2020 gas chromatograph (Shimadzu, Kyoto, Japan) equipped with a thermal conductivity detector (TCD), according to the analytical procedure described previously (Chmielarczyk et al., 2025). Briefly, qualitative analysis was performed using MS line equipped with HP-PLOT/U capillary column (Agilent Technologies), whereas hydrogen concentrations were quantified using TCD line equipped with a 5 Å molecular sieve-packed column.. Quantitative analyses were based on calibrations based on certified gas standards (Air Liquide Polska), and the concentrations of individual gaseous metabolites were calculated based on ares of peaks corresponding to particular compounds.

### 2.4. Endotoxin Assay

To exclude the possibility that the observed biological effects resulted from endotoxin contamination rather than bacterial gaseous metabolites, endotoxin levels were determined in the aqueous solution obtained after dissolving the collected gaseous metabolites in 10 mL of endotoxin-free water. Endotoxin detection was performed using the Thermo Scientific™ Pierce™ Rapid Gel Clot Endotoxin Assay Kit based on the Limulus Amebocyte Lysate (LAL) assay (Thermo Fisher Scientific, Rockford, IL, USA), according to the manufacturer’s instructions. Results were interpreted visually based on gel-clot formation. Samples producing no gel clot were considered endotoxin-negative (<0.125 EU/mL), whereas gel formation indicated endotoxin concentrations equal to or above the assay sensitivity (0.125 EU/mL).

### 2.5. Measurement of gas pressure within the bacterial culture system

To compare the effects of individual primary and secondary bile acids on bacterial gaseous metabolite production, the pressure generated within the closed bacterial culture system was monitored throughout incubation. Gas pressure was measured using a Benetech GM520 digital pressure manometer (maximum pressure, 150 kPa) connected directly to the valve of the 2-L culture flask described in Section 2.2. Measurements were performed after 0, 24, 36, 48, and 72 h of incubation and reflected the total pressure generated by gaseous metabolites accumulating within the closed culture system during bacterial growth.

No pressure measurements were performed for cultures supplemented with chenodeoxycholic acid (CDCA), as no detectable gaseous metabolite production was observed under these conditions. All remaining experimental conditions were analyzed in two independent experiments.

### 2.6. Cell culture

Human colorectal epithelial Caco-2 cells (Sigma-Aldrich, Lot 17H003) and human pancreatic epithelial PANC-1 cells (American Type Culture Collection, ATCC) were cultured according to the suppliers’ recommendations. Caco-2 cells were maintained in Dulbecco’s Modified Eagle Medium (DMEM, Sigma-Aldrich) supplemented with 10% fetal calf serum (FCS), 1% non-essential amino acids, 0.2 mM L-glutamine, and 1% penicillin–streptomycin–neomycin solution. PANC-1 cells were cultured in DMEM supplemented with 10% fetal bovine serum (FBS) and 1% penicillin–streptomycin. Both cell lines were maintained at 37°C in a humidified atmosphere containing 5% CO₂ and routinely passaged at 70–80% confluence using trypsin/EDTA. Cell cultures were regularly monitored microscopically and routinely tested for *Mycoplasma* contamination by PCR.

### 2.7. Exposure of epithelial cells to bacterial gaseous metabolites and assessment of apoptosis and necrosis

To evaluate the biological activity of bacterial gaseous metabolites generated under different primary and secondary bile acid conditions, Caco-2 and PANC-1 cells were seeded onto sterile glass coverslips placed in six-well culture plates and cultured until approximately 90% confluence was reached. The culture plates were subsequently transferred into a sealed gas exposure chamber, which was connected to the bacterial gas production system described in Section 2.2 (Figure 2). This custom-designed exposure system enabled the continuous transfer of freshly generated bacterial gaseous metabolites from the bacterial culture flasks to the cell culture chamber throughout the 48-h incubation period while maintaining standard cell culture conditions.

**Figure 2.**
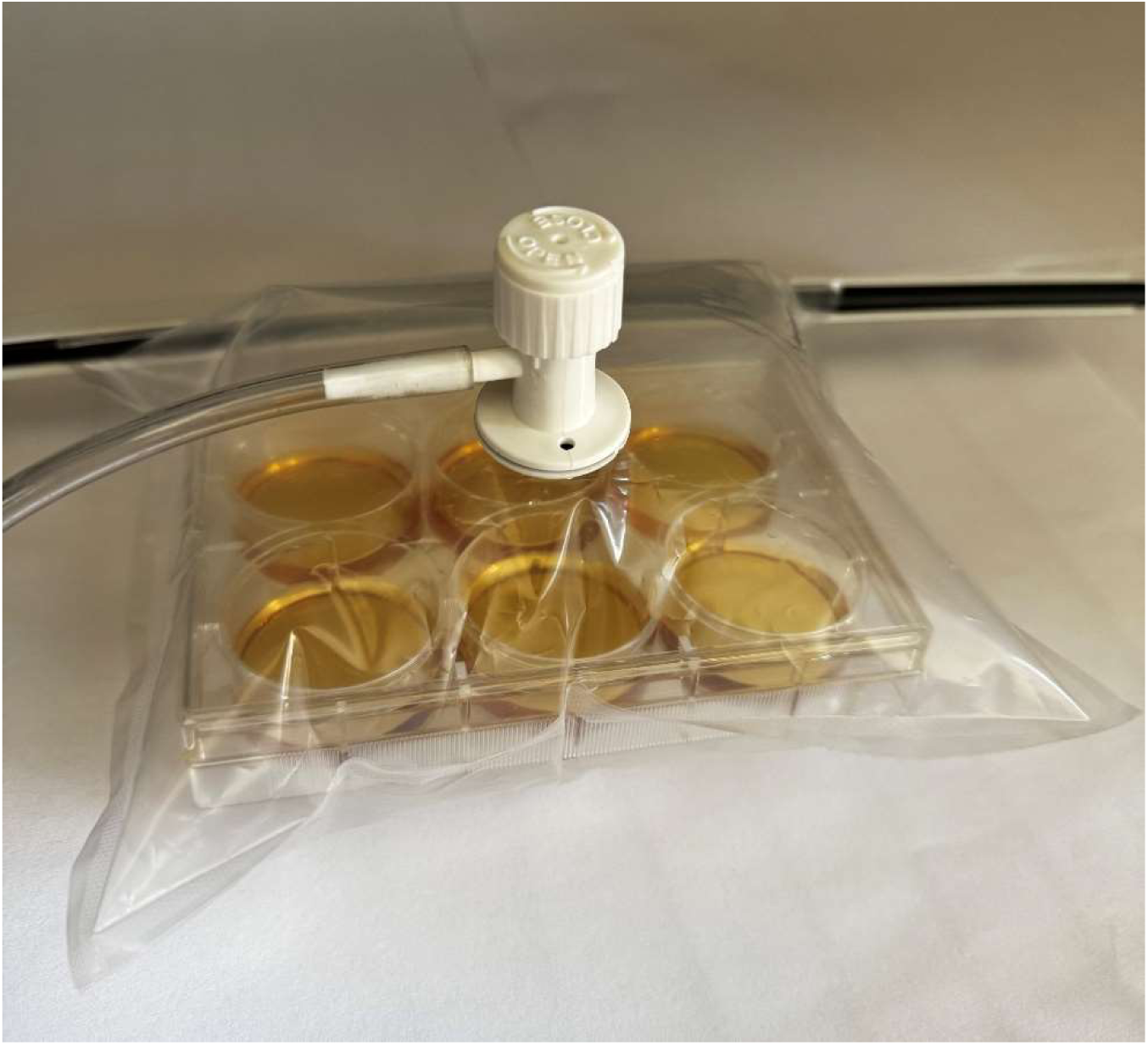
Experimental setup used for indirect exposure of epithelial cell cultures to bacterial gaseous metabolites. Cell culture plates were incubated in sealed chambers connected to bacterial culture flasks generating gaseous metabolites.

Following exposure, apoptosis and necrosis were assessed using the Annexin V-FLUOS Apoptosis Detection Kit (Roche Diagnostics, Mannheim, Germany) according to the manufacturer’s instructions. Cells were stained with Annexin V-FLUOS and propidium iodide and examined under a fluorescence microscope using DAPI, FITC, and TRITC filter sets at ×20 magnification. The percentages of viable, apoptotic, and necrotic cells were determined for each experimental condition and compared with untreated control cultures.

A representative image of the custom-designed gas exposure chamber used for culturing epithelial cells in an atmosphere of bacterial gaseous metabolites is presented in Figure 2.

### 2.8. RNA isolation, library preparation, sequencing and transcriptomic analysis

Following 48 h exposure to bacterial gaseous metabolites generated under different primary and secondary bile acid conditions, total RNA was isolated from Caco-2 and PANC-1 cells using the Total RNA Mini/Midi/Maxi Kit (A&A Biotechnology) according to the manufacturer’s instructions, including an additional genomic DNA removal step. RNA quality and integrity were assessed prior to library preparation. Poly(A)+ mRNA was enriched using the NEBNext® Poly(A) mRNA Magnetic Isolation Module (New England Biolabs), and strand-specific cDNA libraries were prepared using the NEBNext® Ultra™ II Directional RNA Library Prep Kit for Illumina® (New England Biolabs). Library quality was verified using an Agilent 2100 Bioanalyzer with High Sensitivity DNA chips, and library concentrations were determined by quantitative PCR.

High-throughput sequencing was performed on an Illumina NovaSeq platform in paired-end 150 bp (PE150) mode, generating approximately 9 GB of sequencing data per sample. Raw sequencing reads were processed using a standard RNA-seq bioinformatics workflow, including quality assessment, alignment to the human reference genome, transcript quantification, differential gene expression analysis, and functional enrichment based on Gene Ontology (GO) and Kyoto Encyclopedia of Genes and Genomes (KEGG) pathway analyses.

To elucidate the biological effects of bacterial gaseous metabolites, transcriptomic profiles of Caco-2 and PANC-1 cells exposed to gaseous metabolites generated in the presence of individual primary and secondary bile acids were compared with those of untreated control cells. Particular attention was focused on signaling pathways associated with epithelial barrier integrity, inflammatory responses, oxidative stress, apoptosis, cellular adaptation to stress, extracellular matrix remodeling, and carcinogenesis. Functional interpretation of differentially expressed genes was performed using current pathway annotation resources and recent transcriptomic literature (Tomusiak-Plebanek et al., 2025).

## 3. Results

### 3.1. Primary and secondary bile acids differentially regulate the gas-producing phenotype of *Escherichia coli* 33

To determine whether individual bile acids modulate bacterial gaseous metabolism, the high gas-producing Escherichia coli EC33 strain was cultured in the presence of selected primary and secondary bile acids. Visual assessment of extracellular gas accumulation after 48 h revealed pronounced bile acid-dependent differences in the gas-producing phenotype of EC33 (Table 1). Complete filling of the Tedlar® gas collection bags (+++) was consistently observed in cultures supplemented with cholic acid (CA) and deoxycholic acid (DCA), indicating the highest intensity of gaseous metabolite production. In contrast, chenodeoxycholic acid (CDCA) completely inhibited visible gas accumulation, whereas ursodeoxycholic acid (UDCA) produced only trace amounts of gas. Lithocholic acid (LCA) induced an intermediate phenotype characterized by partial gas accumulation.

These qualitative observations were confirmed by quantitative pressure measurements (Figure 3), which demonstrated marked differences in both the kinetics and intensity of bacterial gaseous metabolite production depending on the bile acid present in the culture medium. The highest headspace pressure values were recorded in cultures supplemented with DCA, followed by CA, whereas LCA induced only moderate pressure increases and UDCA produced minimal pressure changes. No measurable increase in headspace pressure was observed in cultures supplemented with CDCA, confirming complete suppression of bacterial gaseous metabolite production under these conditions.

**Figure 3.**
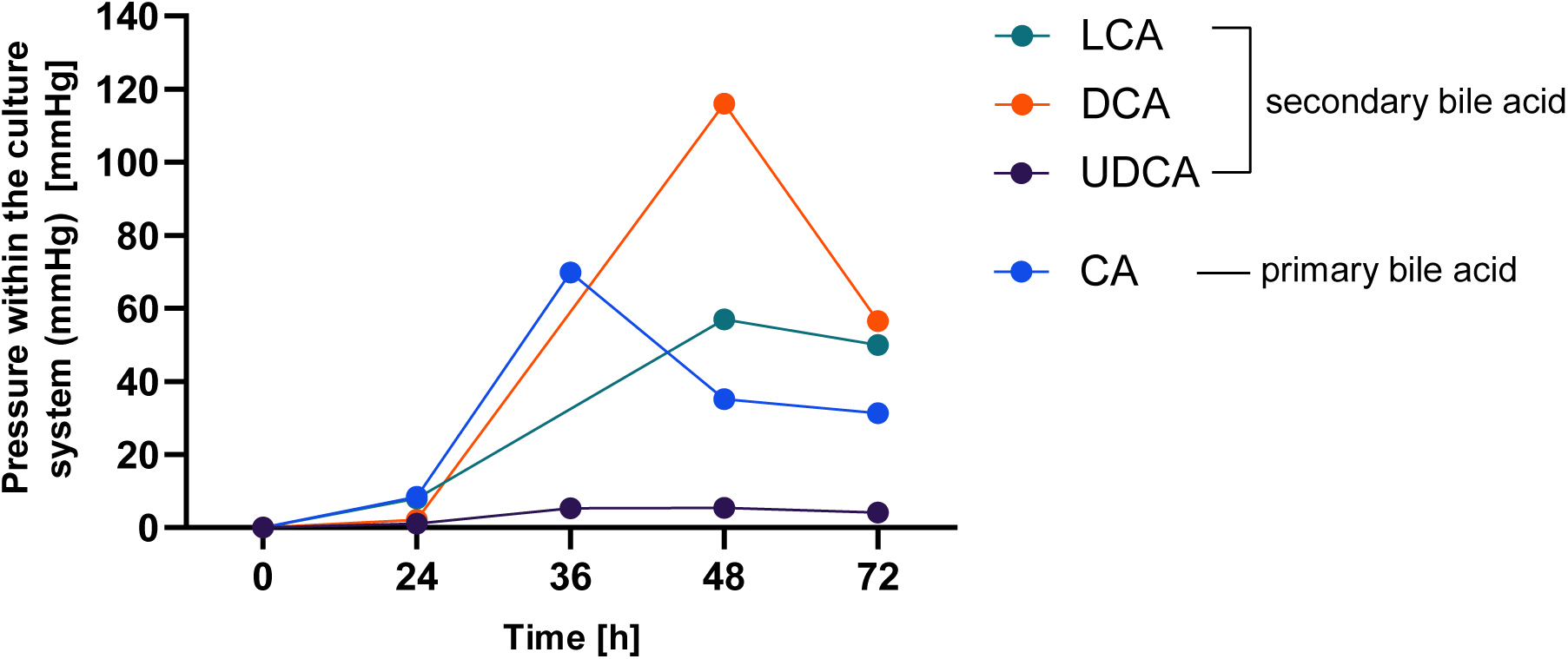
Time-course of pressure generated within the closed bacterial culture system during growth of the high gas-producing EC33 strain in the presence of individual primary and secondary bile acids.

Quantitative GC analysis further demonstrated that individual bile acids significantly influenced both the quantity and composition of gaseous metabolites produced by *E. coli* EC33 (Table 2). Cholic acid (CA) consistently induced the highest total gas production, exceeding 1 L after 48 h of incubation, whereas DCA generated a comparable total gas volume but exhibited a markedly lower proportion of hydrogen, indicating a distinct gaseous metabolic profile. In contrast, UDCA resulted in substantially lower gas production, while gas generation in the presence of LCA showed considerable variability between experimental replicates, suggesting that this bile acid induces a less stable metabolic response. The observed pressure profiles (Figure 3) indicate initial gas accumulation, leading to a gradual increase in pressure inside the sampling bags. At longer incubation times, a stabilization of pressure could be expected as the metabolic activity of the bacteria decreased owing to substrate depletion, product accumulation, or increasing pressure within the system. However, instead of reaching a stable plateau, the pressure subsequently decreased. There are a few processes that may be responsible for this effect. First, part of the generated CO_2_ may be dissolved in the culture medium and subsequently participate in carbonate-bicarbonate equilibria, thereby decreasing the amount of gas present in the bag. Changes in the metabolic activity of E. coli during prolonged cultivation may also have altered the balance between gas production and consumption, including the possible utilization of previously generated H_2_ or CO_2_. Nevertheless, the most likely explanation is associated with the mechanical properties of the flexible sampling bags. During filling, the folds of the bag gradually unfold, and the polymer walls undergo expansion and relaxation. Therefore, if the increase in bag volume becomes greater than the relative increase in the amount of gas, the measured pressure may decrease even though gas production is still occurring. It should be underlined that, because the bags were flexible and extensible, the pressure kinetics cannot be interpreted solely in terms of bacterial metabolic activity without an independent pressure– volume calibration; that is why the analyses were supplemented by the measurements of volume of produced gas and its quantitative and qualitative GC-MS/TCD analyses.

**Table 2.**
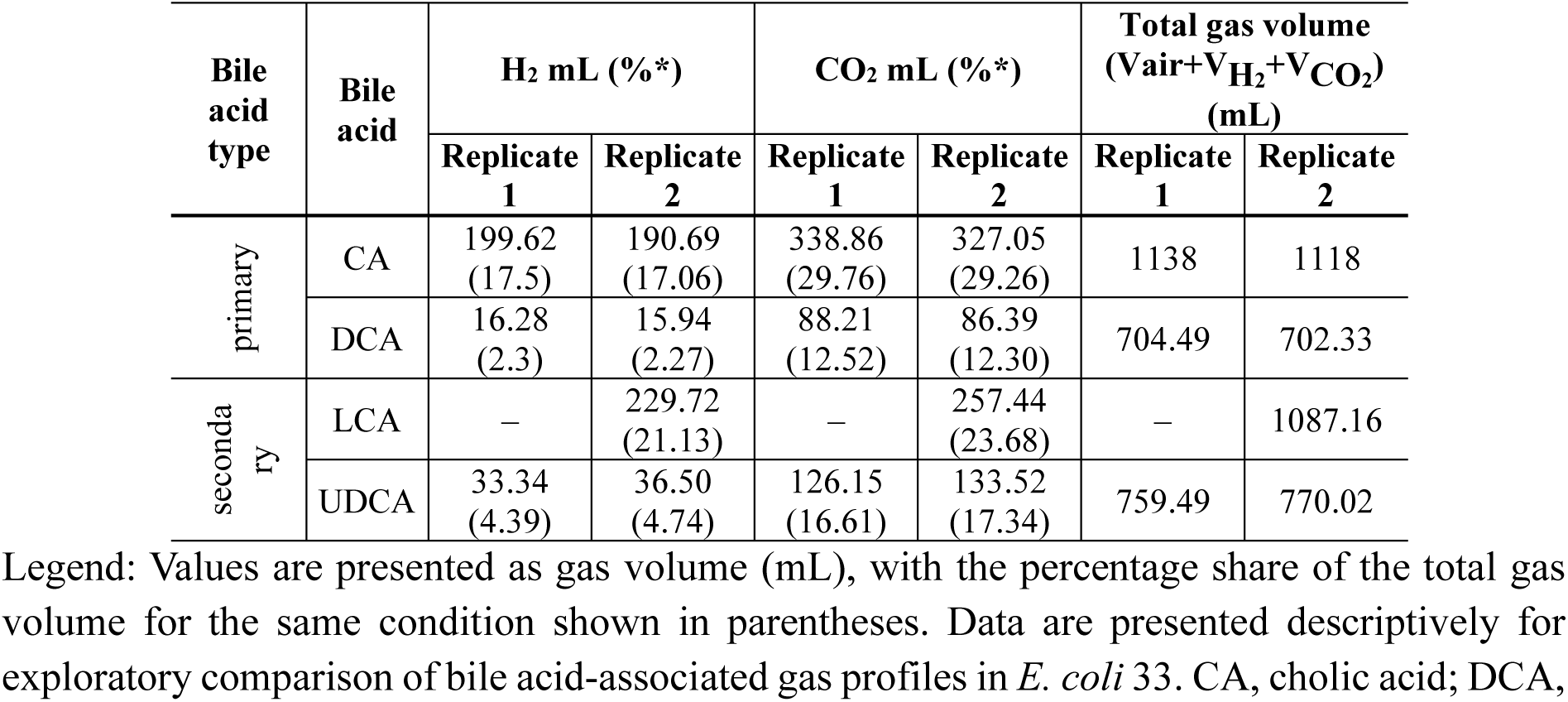
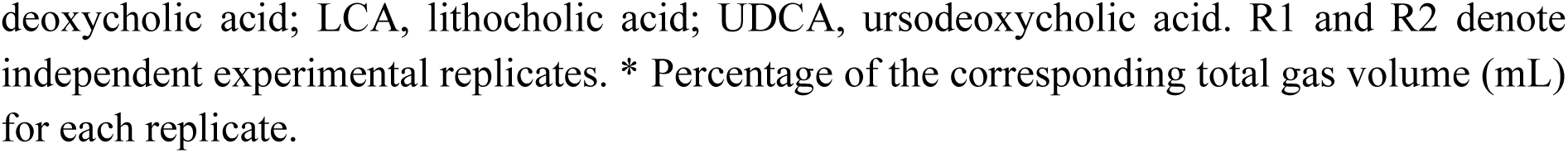
Quantitative production of gaseous metabolites determined for *E. coli* 33 strain after 48 h of incubation on MacConkey agar with or without primary or secondary bile acids.

Regardless of the bile acid tested, carbon dioxide represented the predominant gaseous metabolite, whereas hydrogen production varied markedly depending on bile acid composition. Collectively, these findings demonstrate that individual primary and secondary bile acids regulate not only the intensity of bacterial gaseous metabolite production but also the metabolic signature of the gases produced, supporting the concept that changes in bile acid composition can reshape bacterial fermentation pathways and the gas-producing phenotype of intestinal bacteria.

To exclude the possibility that the biological effects observed in subsequent cell culture experiments resulted from endotoxin contamination rather than bacterial gaseous metabolites, all gas samples were analyzed using the Limulus Amebocyte Lysate (LAL) assay. No detectable endotoxin contamination was observed in any sample at the assay sensitivity threshold of 0.125 EU/mL, confirming that the collected gaseous metabolite preparations were suitable for subsequent biological analyses. Having established bile acid-dependent differences in bacterial gaseous metabolite production and verified the purity of the collected gas samples, we next investigated whether these metabolites exert direct cytotoxic effects on intestinal and pancreatic epithelial cells.

### 3.2. Bacterial gaseous metabolites induce only limited apoptosis and necrosis in epithelial cells

Based on the gas production experiments, CA and DCA were selected for subsequent biological analyses because they consistently induced the highest production of bacterial gaseous metabolites.

To determine whether bacterial gaseous metabolites exert direct cytotoxic effects, Caco-2 intestinal epithelial cells and PANC-1 pancreatic epithelial cells were exposed for 48 h to gaseous metabolites generated by the high gas-producing E. coli EC33 strain cultured under different bile acid conditions. Cell death was subsequently evaluated by Annexin V/propidium iodide staining (Table 3).

**Table 3.** Effect of bacterial gaseous metabolites generated by Escherichia coli EC33 cultured in the presence of cholic acid (CA) or deoxycholic acid (DCA) on epithelial cell viability, apoptosis, and necrosis after 48 h of exposure. Results are expressed as percentages of total cells.

| Experimental condition | PANC-1 cells |  |  | Caco-2 cells |  |  |
| --- | --- | --- | --- | --- | --- | --- |
|  | Viability (%) | Apoptosis (%) | Necrosis (%) | Viability (%) | Apoptosis (%) | Necrosis (%) |
| CA | 92 | 3 | 5 | 92 | 0 | 8 |
| DCA | 95 | 2 | 3 | 91 | 0 | 9 |
| Control | 98 | 0 | 2 | 98 | 0 | 2 |

Legend: CA - cholic acid; DCA - deoxycholic acid. Control refers to cells cultured under the same experimental conditions without exposure to bacterial gaseous metabolites.

Overall, exposure to bacterial gaseous metabolites induced only minimal cytotoxicity. In both epithelial cell lines, cell viability remained above 90% under all experimental conditions. No increase in apoptotic cells was observed in Caco-2 cultures, whereas PANC-1 cells exhibited only a slight increase in apoptosis following exposure to gaseous metabolites generated in the presence of deoxycholic acid (DCA). Similarly, necrotic cell death remained low in both cell lines and differed only marginally from the corresponding control cultures.

These findings indicate that bacterial gaseous metabolites, under the experimental conditions used, do not induce extensive epithelial cell death. Instead, they appear to trigger subtle cellular responses while largely preserving cell viability. Importantly, these observations demonstrate that the experimental exposure model itself does not exert nonspecific cytotoxic effects and therefore provides an appropriate platform for investigating early molecular responses to bacterial gaseous metabolites.

The absence of substantial apoptosis and necrosis is particularly relevant for the interpretation of the subsequent transcriptomic analyses. Since epithelial cell viability remained largely preserved, the observed changes in gene expression are unlikely to represent secondary consequences of cell death. Rather, they most likely reflect specific biological responses induced by bacterial gaseous metabolites, supporting the hypothesis that these compounds function primarily as signaling molecules capable of modulating epithelial cell physiology rather than as direct cytotoxic agents.

### 3.3. Transcriptomic remodeling induced by bacterial gaseous metabolites

#### 3.3.1. Caco-2 cells exposed to bacterial gaseous metabolites generated in the presence of cholic acid (CA)

RNA sequencing revealed that exposure of Caco-2 intestinal epithelial cells to bacterial gaseous metabolites generated by E. coli EC33 cultured in the presence of cholic acid (CA) resulted in extensive transcriptional remodeling (Table 4). Differentially expressed genes (DEGs) were primarily associated with inflammatory signaling, cellular stress responses, extracellular matrix organization, metabolic adaptation, and regulation of epithelial homeostasis.

**Table 4.**
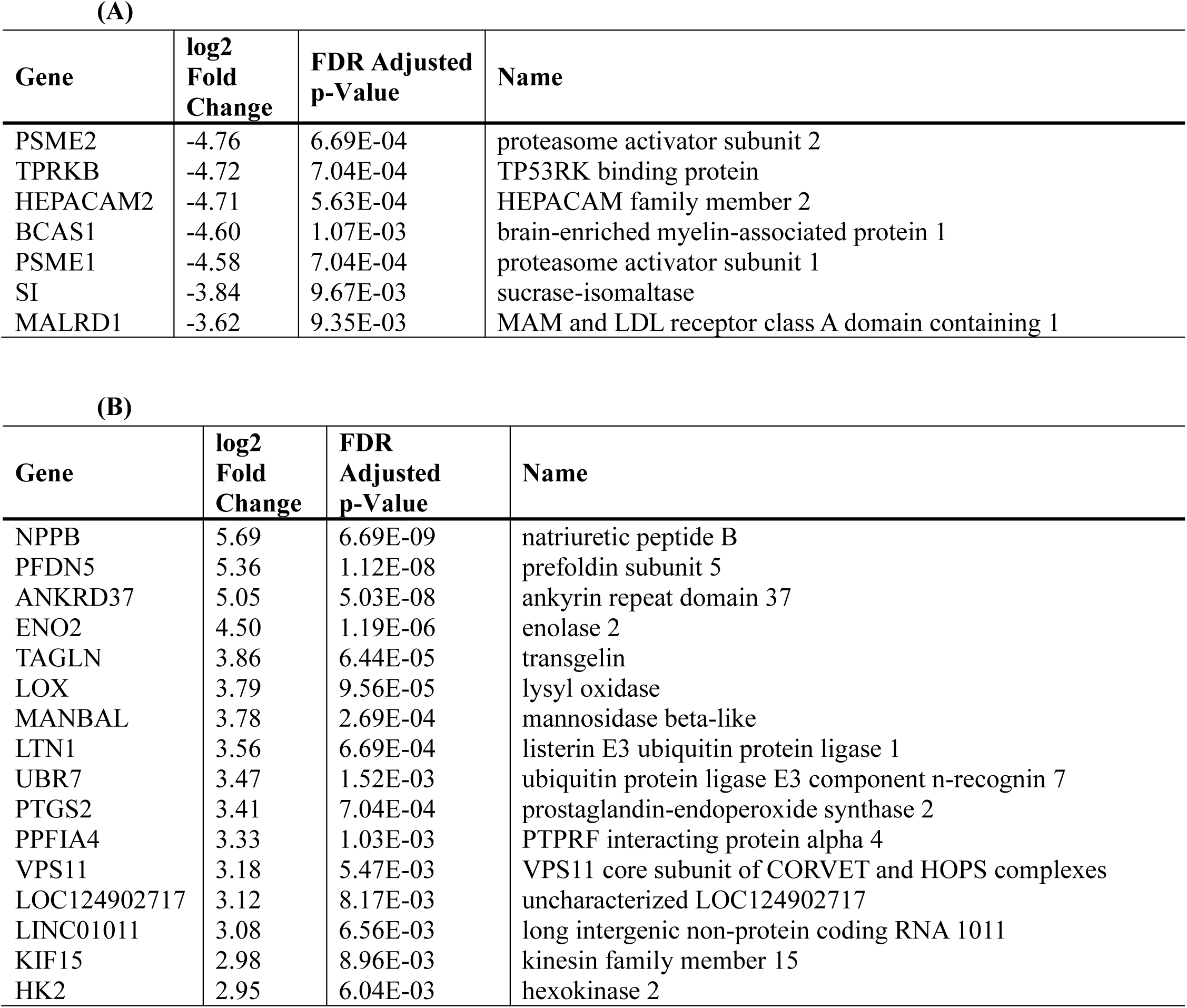
Significantly **(A)** downregulated and **(B)** upregulated differentially expressed genes (DEGs) in the Caco-2 cell line exposed to gas pressure generated by *Escherichia coli* 33, in culture medium supplemented with cholic acid (CA)

Among the most strongly downregulated genes, TPRKB exhibited a marked decrease in expression (log₂FC = −4.72). TPRKB encodes a regulatory component of the TP53RK complex involved in p53-associated signaling and cellular stress responses. Alterations in pathways related to p53 regulation have been implicated in genomic instability and cancer progression; however, the biological consequences of TPRKB downregulation in response to bacterial gaseous metabolites require further functional validation (Goswami et al. 2019).

In contrast, several genes associated with inflammatory activation and cellular adaptation were strongly upregulated. The most prominent change was observed for PTGS2 (COX-2) (log₂FC = +3.41; FDR = 7.04 × 10⁻⁴). COX-2 is a key mediator of prostaglandin synthesis, particularly prostaglandin E₂ (PGE₂), and has been extensively linked to inflammatory signaling, epithelial proliferation, angiogenic responses, and colorectal cancer progression through pathways involving WNT/β-catenin, PI3K/AKT, MAPK, and NF-κB signaling (Oshima et al. 1996; Greenhough et al. 2009; Wang and Dubois 2010). Therefore, the strong induction of PTGS2 suggests that CA-derived bacterial gaseous metabolites may promote a transcriptional state characterized by enhanced inflammatory and epithelial stress signaling.

#### 3.3.2. PANC-1 cells exposed to bacterial gaseous metabolites generated in the presence of cholic acid (CA)

Compared with Caco-2 cells, PANC-1 cells exhibited a substantially more restricted transcriptional response following exposure to CA-derived bacterial gaseous metabolites (Table 5). Differentially expressed genes were mainly associated with oxidative stress adaptation, autophagy-related processes, vesicular trafficking, and cytoskeletal organization.

**Table 5.**
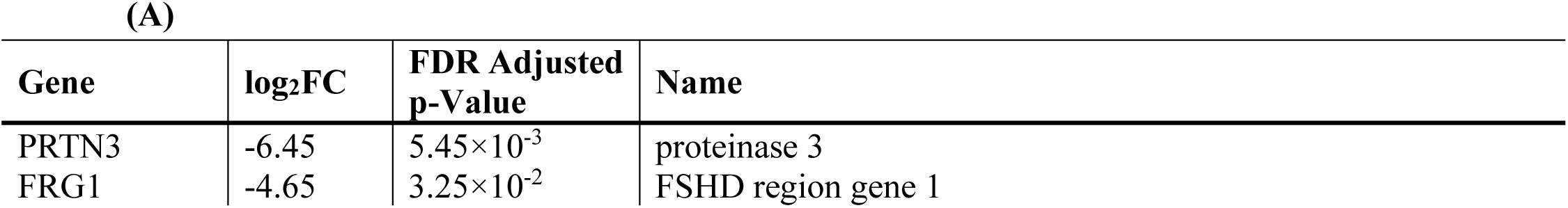
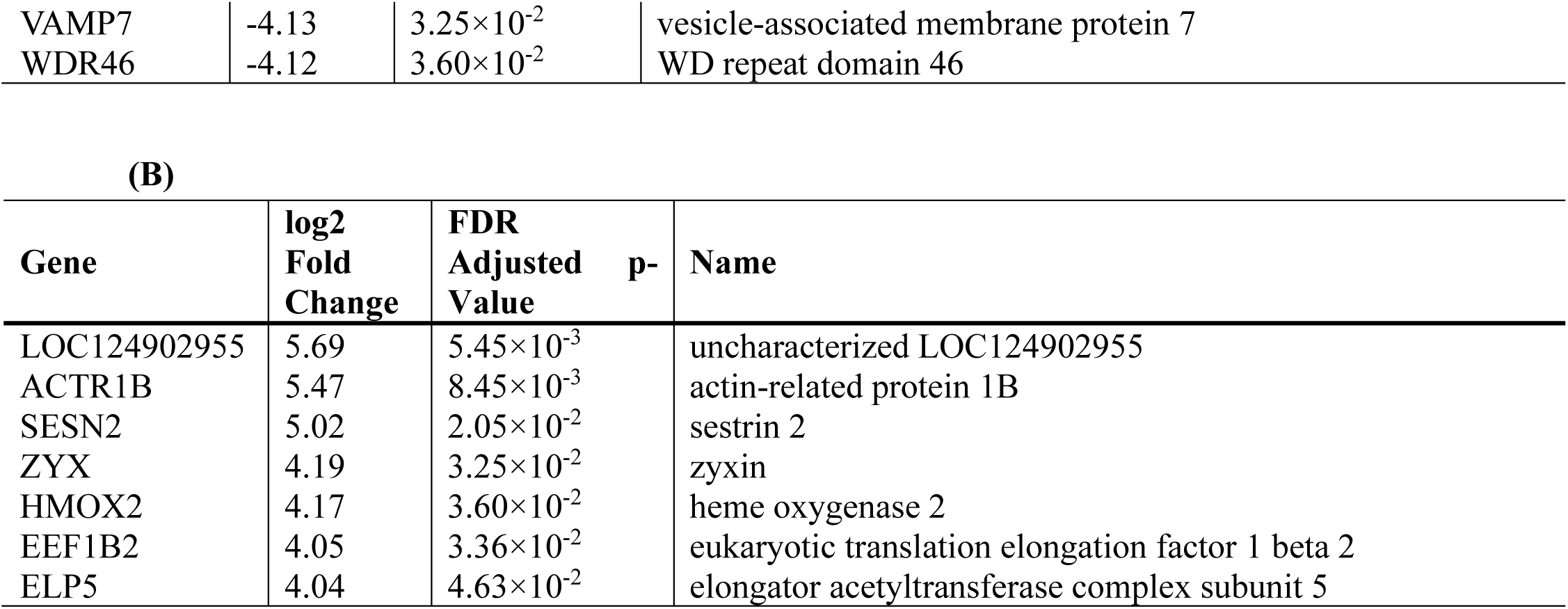
Significantly **(A)** downregulated and **(B)** upregulated differentially expressed genes (DEGs) in the PANC1 cell line exposed to gas pressure generated by *Escherichia coli* 33, in culture medium supplemented with cholic acid (CA)

The most strongly downregulated gene was VAMP7 (log_2_FC = −4.13; FDR = 3.25 10^-2^), a vesicle-associated protein involved in intracellular trafficking, membrane fusion, and autophagic processes. VAMP7 has been implicated in cancer cell migration, invasion, and survival mechanisms, including autophagy-dependent adaptation to cellular stress (Aoyagi et al. 2018; Russell and Guan 2022). Although the functional significance of VAMP7 downregulation in this experimental model remains to be established, its altered expression suggests modulation of intracellular stress-response pathways.

Conversely, genes involved in oxidative stress regulation and cellular adaptation, including SESN2, HMOX2, and ZYX (zyxin), were upregulated. SESN2 and HMOX2 contribute to redox homeostasis and cellular stress adaptation, whereas ZYX participates in cytoskeletal organization and focal adhesion signaling. Zyxin directly promotes cell proliferation, migration, invasion, and metastatic potential through activation of AKT/mTOR signaling and focal adhesion–dependent cytoskeletal remodeling. Its strong upregulation (log_2_FC = 4.19) suggests a central functional role in driving the malignant phenotype of pancreatic cancer cells.

The observed transcriptional profile indicates that PANC-1 cells respond to CA-derived gaseous metabolites primarily through activation of adaptive stress pathways rather than broad transcriptional reprogramming. (Zhong et al. 2019; Cai et al. 2023).

#### 3.3.3. Caco-2 cells exposed to bacterial gaseous metabolites generated in the presence of deoxycholic acid (DCA)

Exposure of Caco-2 cells to bacterial gaseous metabolites generated under deoxycholic acid (DCA) conditions resulted in a markedly stronger transcriptional response compared with CA- derived metabolites (Table 6). The regulated genes represented a broad transcriptional signature involving inflammatory signaling, extracellular matrix remodeling, epithelial plasticity, metabolic adaptation, and pathways previously associated with colorectal cancer progression. Among the most strongly induced genes were NR4A2, PIM1, JUN, FOS, MMP1, PLAUR, CEACAM5, and CEACAM6. PIM1 is a serine/threonine kinase involved in cell survival, proliferation, metabolic regulation, and resistance to stress-induced apoptosis (Choudhury et al. 2024). JUN and FOS represent components of the AP-1 transcription factor complex, which regulates inflammatory responses, proliferation, and cellular plasticity. Similarly, MMP1 and PLAUR are associated with extracellular matrix degradation, invasion, and tissue remodeling (Conlon and Murray 2019).

**Table 6.**
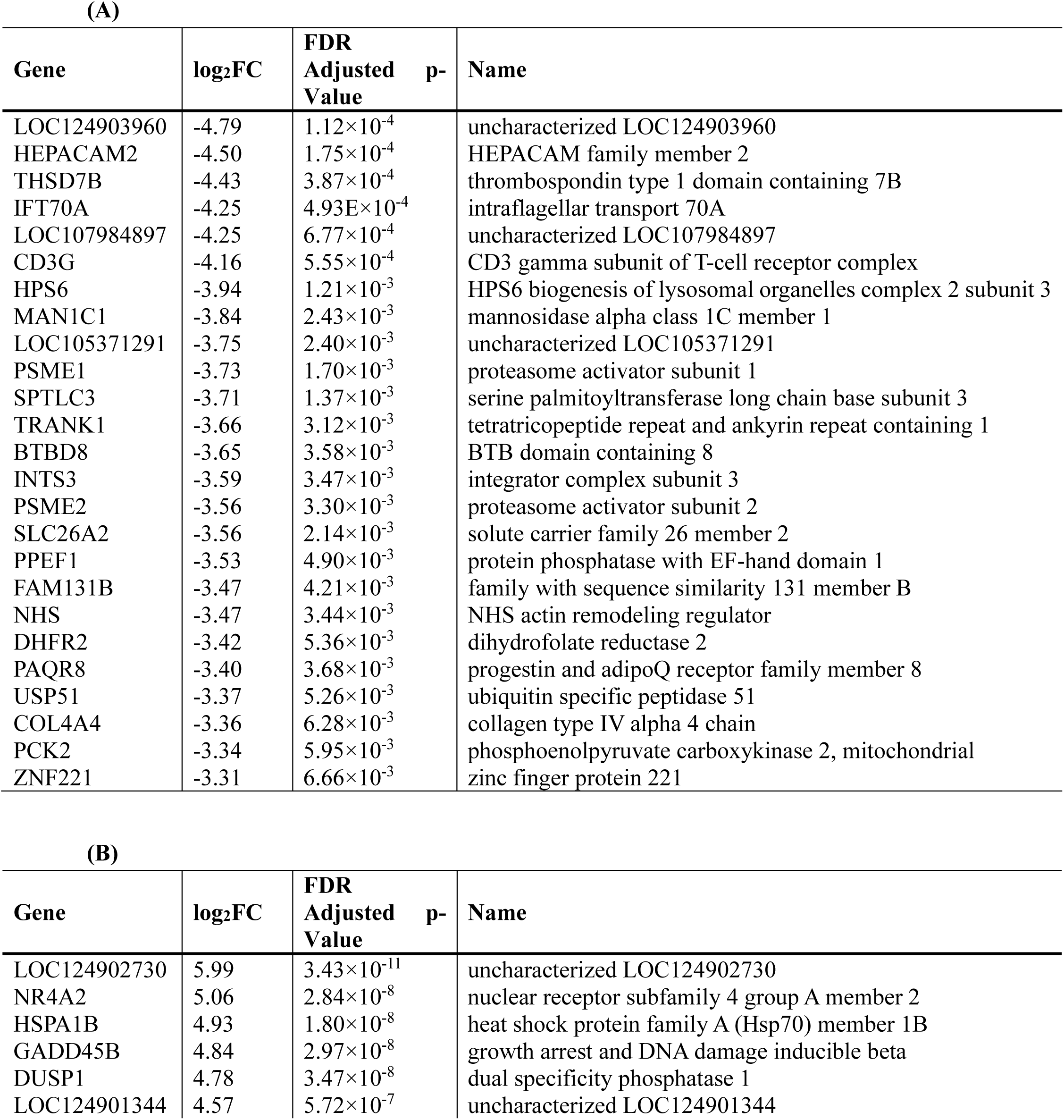
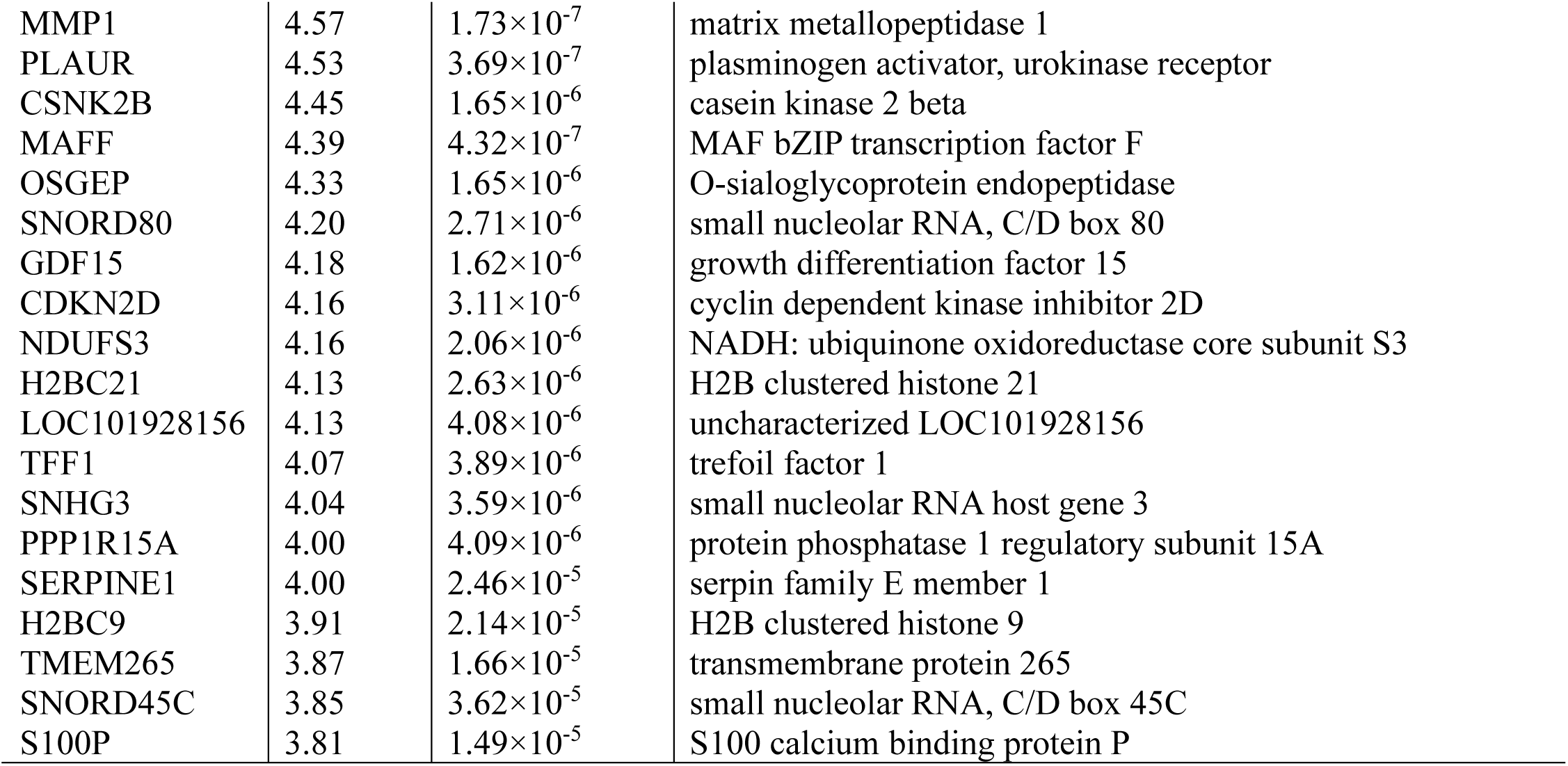
Significantly **(A)** downregulated and **(B)** upregulated differentially expressed genes (DEGs) in the Caco-2 cell line exposed to gas pressure generated by *Escherichia coli* 33, in culture medium supplemented with deoxycholic acid (DCA)

The induction of CEACAM5 and CEACAM6, which are widely recognized biomarkers associated with colorectal cancer biology, further supports the presence of a transcriptional phenotype characterized by altered epithelial plasticity and cancer-associated signaling (Aldilaijan et al. 2023; Liang et al. 2024). In parallel, downregulation of genes involved in proteasome regulation and epithelial organization, including PSME1, PSME2, and HEPACAM2, suggests additional alterations in cellular homeostasis(Xie et al. 2025; Wang et al. 2025). Collectively, these findings indicate that DCA-derived bacterial gaseous metabolites exert a substantially stronger transcriptional effect on intestinal epithelial cells than CA-derivedvmetabolites, promoting a gene expression profile enriched in inflammatory, remodeling, and stress-associated pathways.

#### 3.3.4. PANC-1 cells exposed to bacterial gaseous metabolites generated in the presence of deoxycholic acid

Following exposure to DCA-derived bacterial gaseous metabolites, PANC-1 cells displayed considerably fewer transcriptional changes than Caco-2 cells (Table 7). The identified DEGs were mainly associated with proteostasis, protein degradation, and cytoskeletal organization. The most prominent alterations included upregulation of DCAF11 and PSME2 together with downregulation of FRG1. PSME2 encodes a proteasome activator subunit involved in regulation of proteasomal activity and cellular adaptation to stress conditions, whereas DCAF11 participates in ubiquitin-dependent protein degradation pathways (Guo et al. 2021). FRG1 contributes to cytoskeletal organization and cellular structural stability, and its altered expression has been associated with changes in cellular plasticity in cancer models (Khan et al. 2021).

**Table 7.** Significantly **(A)** downregulated and **(B)** upregulated differentially expressed genes (DEGs) in the PANC1 cell line exposed to gas pressure generated by *Escherichia coli* 33, in culture medium supplemented with deoxycholic acid (DCA)

| Gene | log<sub>2</sub>FC | FDR<br>Adjusted<br>p-Value | Name |
| --- | --- | --- | --- |
| FRG1 | -4.78 | 4.77E-02 | FSHD region gene 1 |

(B)
| Gene | log<sub>2</sub>FC | FDR<br>Adjusted<br>p-Value | Name |
| --- | --- | --- | --- |
| DCAF11 | 5.42 | 1.70E-03 | proteasome activator subunit 2 |
| PSME2 | 4.44 | 4.77E-02 | DDB1 and CUL4 associated factor 11 |

Although the magnitude of transcriptomic remodeling was limited compared with intestinal epithelial cells, these findings demonstrate that pancreatic epithelial cells remain molecularly responsive to bacterial gaseous metabolites, primarily through selective regulation of stress- adaptation and proteostasis pathways.

#### 3.3.5. KEGG pathway enrichment analysis

KEGG pathway enrichment analysis demonstrated that the biological effects of bacterial gaseous metabolites were strongly influenced by both the epithelial cell type and the bile acid environment in which these metabolites were generated (Figure 4).

**Figure 4.**
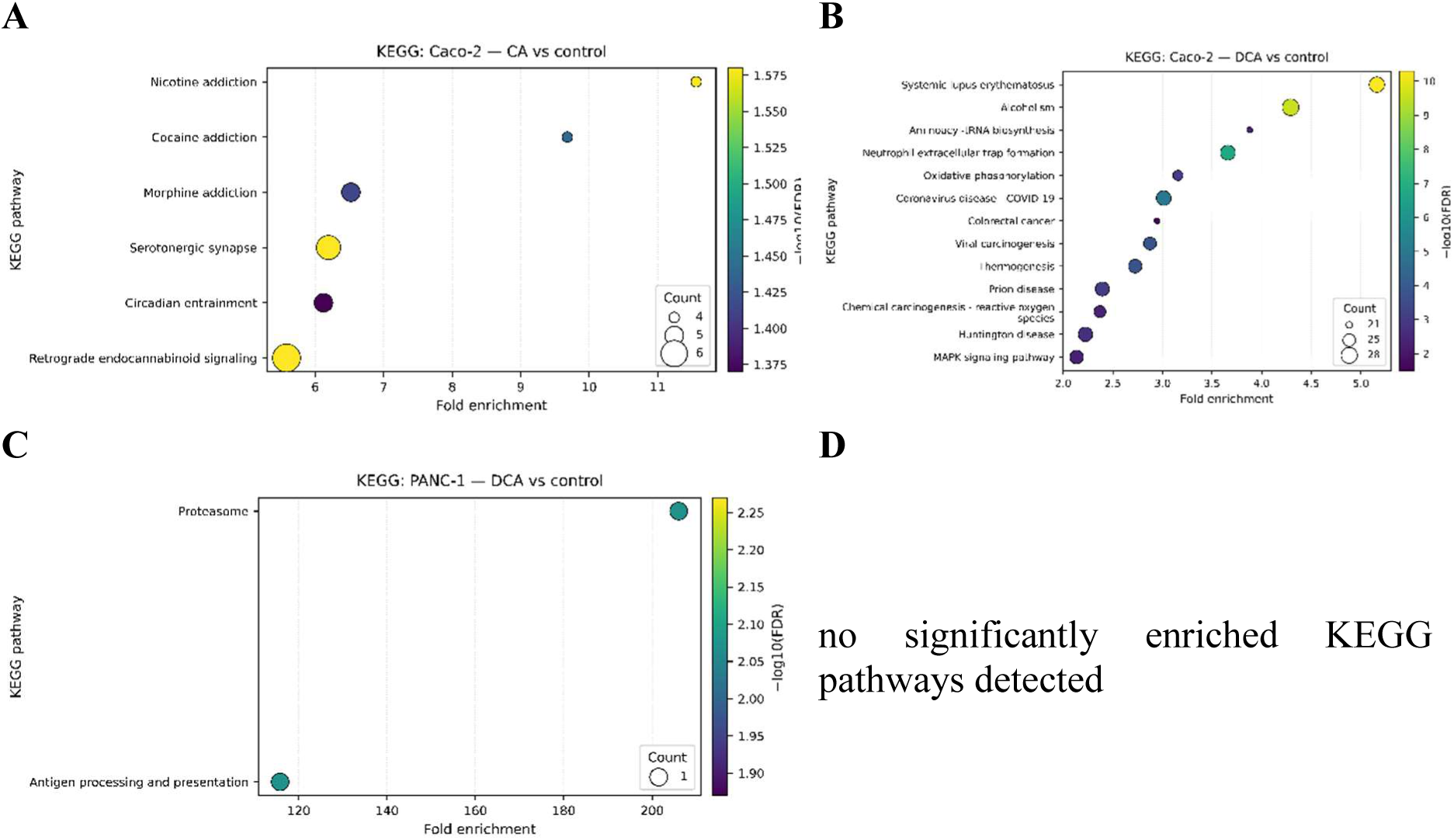
KEGG pathway enrichment analysis. Bubble plots show enriched KEGG pathways for differentially expressed genes (DEGs). (A) Caco-2: CA vs control; (B) Caco-2: DCA vs control; (C) PANC-1: DCA vs control; (D) PANC-1: CA vs control (no significantly enriched KEGG pathways detected). The x-axis indicates fold enrichment. Bubble size represents the number of DEGs mapped to each pathway (Count). Bubble color indicates significance as −log_10_(FDR) (Benjamini–Hochberg adjusted p-value). Up to the top 15 pathways are displayed per panel (fewer when fewer significant terms are available).

**Figure 5.**
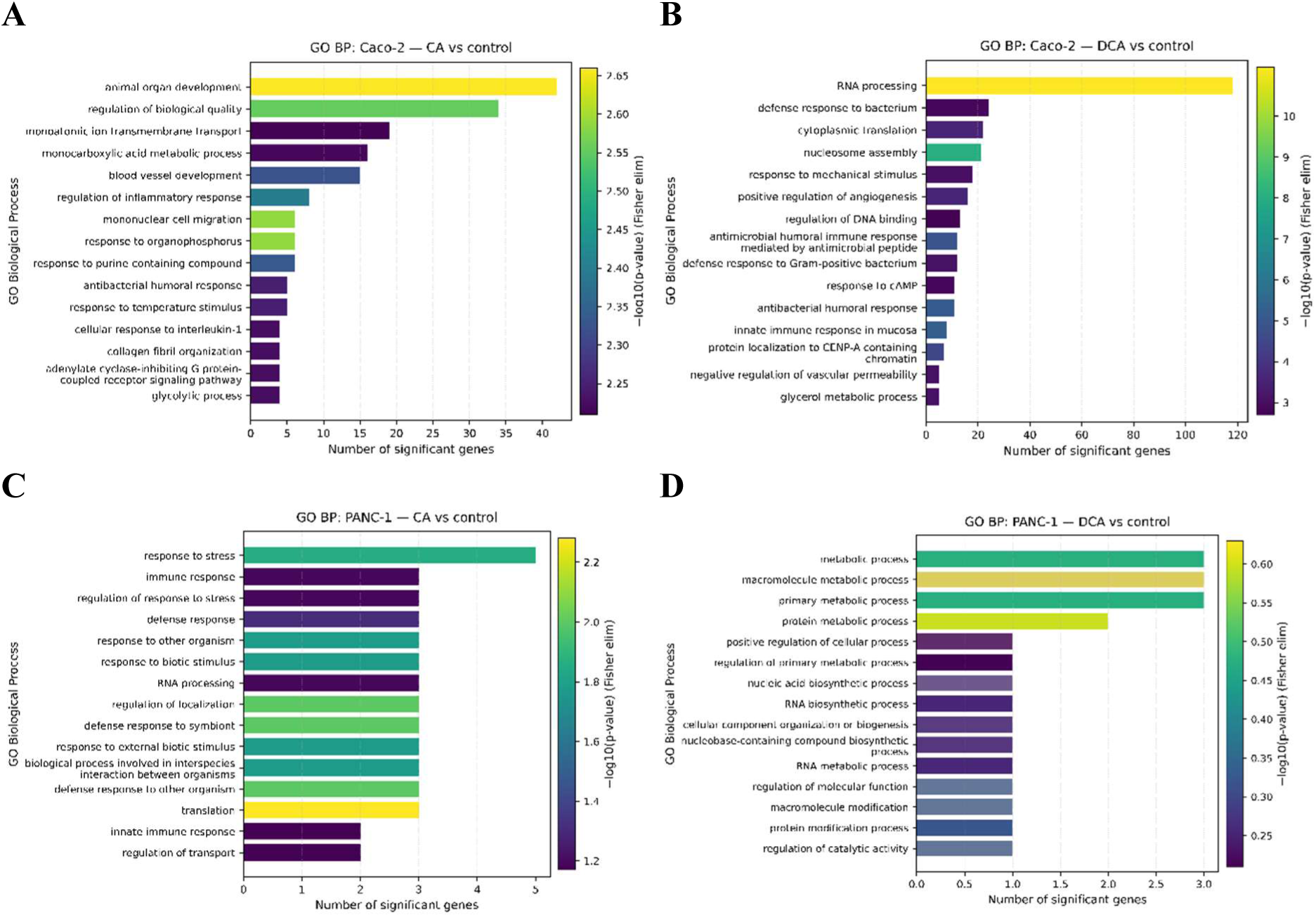
GO Biological Process enrichment. Bar plots show enriched Gene Ontology (GO) Biological Process terms for DEGs. (A) Caco-2: CA vs control; (B) Caco-2: DCA vs control; (C) PANC-1: CA vs control; (D) PANC-1: DCA vs control. Bar length (x-axis) indicates the number of significantly annotated DEGs (Significant) assigned to each GO term. Bar color indicates significance as −log10(p-value) from Fisher’s exact test with the elim algorithm (Fisher elim). Up to the top 15 terms are displayed per panel (fewer when fewer terms are available).

The strongest pathway enrichment was observed in Caco-2 cells exposed to DCA-derived metabolites. Significantly enriched pathways included inflammatory signaling, extracellular matrix remodeling, cellular stress responses, regulation of proliferation, and cancer-associated pathways. These findings are consistent with the established role of chronic inflammatory signaling and extracellular matrix remodeling in intestinal epithelial dysfunction and colorectal cancer progression (Candela et al. 2014; Hanahan 2022).

CA-derived metabolites also induced pathway enrichment in Caco-2 cells, although the response was quantitatively weaker and primarily associated with adaptive metabolic and stress-response pathways. In contrast, PANC-1 cells displayed substantially fewer enriched pathways, and no significant KEGG enrichment was detected following exposure to CA- derived metabolites despite the presence of individual DEGs.

Together, these data demonstrate that intestinal epithelial cells are substantially more responsive to bacterial gaseous metabolites than pancreatic epithelial cells. Moreover, the distinct transcriptional profiles induced by CA- and DCA-derived metabolites indicate that the biological activity of microbiome-derived gaseous compounds is determined not only by the target cell type but also by the metabolic context in which these compounds are generated. These findings support the concept that bacterial gaseous metabolites represent an additional class of microbiome-derived signaling molecules capable of modulating host cellular pathways in a bile acid-dependent manner.

Exposure to CA-derived gaseous metabolites also resulted in significant pathway enrichment in Caco-2 cells; however, fewer pathways reached statistical significance, indicating activation of a more limited adaptive transcriptional response. In contrast, PANC-1 cells exhibited substantially fewer significantly enriched pathways. Following exposure to DCA-derived bacterial gaseous metabolites, only a limited number of pathways reached statistical significance, whereas no significantly enriched KEGG pathways were identified in PANC-1 cells exposed to CA-derived gases, despite the presence of differentially expressed genes.

Collectively, these results demonstrate that intestinal epithelial cells are considerably more responsive to bacterial gaseous metabolites than pancreatic epithelial cells. Moreover, the markedly stronger transcriptional effects observed following exposure to DCA-derived gases indicate that the biological activity of bacterial gaseous metabolites depends not only on the target cell type but also on the bile acid environment in which these volatile metabolites are generated.

### 3.4. Discussion

The present study demonstrates that bacterial production of gaseous metabolites is a highly strain-dependent phenomenon and may be profoundly modulated by the composition of bile acids. To our knowledge, this is among the first studies combining microbiological, physicochemical, and transcriptomic approaches to investigate bacterial gaseous metabolism in the context of acute pancreatitis. Our findings suggest that specific *Escherichia coli* isolates recovered from a patient with acute pancreatitis possess a unique metabolic phenotype characterized by enhanced gas production under selected bile acid conditions, while the resulting gaseous environment is capable of inducing significant transcriptional responses in both intestinal and pancreatic epithelial cells (Liu et al. 2024; Li et al. 2024; Pan et al. 2024).

One of the major conceptual implications of the present study is the identification of a potential gas-producing microbiome functional phenotype. Unlike conventional bacterial traits, such as antibiotic resistance, virulence, or biofilm formation, the ability to generate biologically active gaseous metabolites has rarely been considered as a functional characteristic of the microbiome. Our findings suggest that this phenotype is determined by the interaction between specific bacterial strains and environmental signals, particularly bile acid composition. Therefore, gas-producing capacity may represent a previously unrecognized functional property of the intestinal microbiota with potential relevance for host physiology and disease susceptibility.

A further important finding of this study is that individual bile acids differentially regulate bacterial gaseous metabolism rather than simply affecting the overall amount of gas produced. Both the volume and composition of gaseous metabolites varied considerably depending on the bile acid present in the culture medium, highlighting the metabolic diversity that may exist even within a single bacterial species. Although strain-specific variability has previously been described for virulence factors, biofilm formation, and metabolic adaptation, gaseous metabolism has received very little attention. Our findings therefore indicate that the production of biologically active gaseous metabolites should be considered a strain-specific functional trait rather than a universal property of E. coli (Touchon et al. 2020; Denamur et al. 2021; Chmielarczyk et al. 2025).

The observed influence of bile acids on bacterial gas production is biologically plausible. Bile acids represent one of the strongest environmental signals encountered by intestinal bacteria and profoundly affect membrane integrity, energy metabolism, stress adaptation, and bacterial gene expression. Our experiments demonstrate that individual bile acids differentially regulate gaseous metabolism, suggesting that bacterial pathways responsible for hydrogen and carbon dioxide production are tightly linked to bile acid sensing mechanisms. Particularly noteworthy was the almost complete inhibition of gas production by chenodeoxycholic acid, whereas cholic acid and deoxycholic acid strongly stimulated gaseous metabolite formation. These findings support the concept that alterations in bile acid composition accompanying biliary disorders and acute pancreatitis may substantially reshape bacterial physiology and consequently modify the metabolic activity of the intestinal microbiota (Fogelson et al. 2023; Malhotra et al. 2023; Pan et al. 2024).

The differences observed in hydrogen and carbon dioxide production further suggest activation of distinct bacterial metabolic pathways under different bile acid conditions. Hydrogen production primarily reflects bacterial fermentative activity and intracellular redox homeostasis, whereas carbon dioxide originates from multiple decarboxylation reactions associated with central carbon metabolism. Consequently, variations in the H_2_/CO_2_ ratio likely reflect global metabolic reprogramming rather than simple quantitative differences in bacterial growth. Future metabolomic studies should clarify which bacterial enzymatic pathways are responsible for these bile acid-dependent alterations in gaseous metabolism.

The pressure measurements performed in parallel with gas composition analyses provide an additional dimension to the biological interpretation of our findings. Although intestinal gases are generally assumed to diffuse freely within the gastrointestinal lumen, localized gas accumulation within poorly ventilated anatomical compartments, including obstructed bile ducts, pancreatic ducts, or necrotic pancreatic collections, may theoretically generate increased local pressure. While our experimental model does not reproduce in vivo conditions directly, the ability of a single bacterial strain to generate measurable pressure supports the hypothesis that bacterial gaseous metabolism may contribute not only to biochemical signaling but also to local mechanical alterations within inflamed tissues. (Pan et al. 2024).

One of the most interesting aspects of the present study is the demonstration that bacterial gaseous metabolites alone, without direct bacterial contact, modify gene expression in epithelial cells. This observation suggests that volatile bacterial products may function as previously underappreciated mediators of host–microbiota communication. Although apoptosis and necrosis remained relatively limited after 48 hours of exposure, transcriptomic analysis revealed activation of multiple signaling pathways associated with inflammation, cellular stress, and carcinogenesis. These molecular alterations probably precede detectable morphological damage and therefore may represent early biomarkers of cellular responses to bacterial gaseous metabolites (Fogelson et al. 2023; Pan et al. 2024).

Among Caco-2 cells exposed to gases generated in the presence of cholic acid, one of the most notable findings was the marked upregulation of *PTGS2* (COX-2), a central mediator of inflammatory responses and one of the best-established drivers of colorectal carcinogenesis. Increased COX-2 expression promotes prostaglandin synthesis, stimulates proliferation, suppresses apoptosis, and enhances angiogenesis through activation of WNT/β-catenin, PI3K/AKT, and MAPK signaling pathways. Simultaneously, the observed downregulation of genes associated with proteasome activity and epithelial differentiation suggests impairment of normal intestinal epithelial homeostasis. Together, these findings indicate that prolonged exposure to bacterial gaseous metabolites may induce a transcriptional program favouring chronic inflammation and epithelial remodeling (Oshima et al. 1996; Greenhough et al. 2009; Wang and Dubois 2010).

The transcriptomic response induced under deoxycholic acid conditions appeared even more pronounced. Upregulation of *MMP1*, *PLAUR*, *NR4A2*, *SERPINE1* and stress-response genes indicates activation of pathways involved in extracellular matrix degradation, epithelial– mesenchymal transition, cellular migration and inflammatory adaptation. Importantly, deoxycholic acid has previously been implicated in colorectal carcinogenesis, and our results raise the possibility that bacterial gaseous metabolites generated in the presence of this bile acid may amplify its biological effects. Whether this interaction contributes to long-term disease progression requires further investigation (Conlon and Murray 2019; Safe and Karki 2021; Aldilaijan et al. 2023; Choudhury et al. 2024).

The response observed in pancreatic PANC-1 cells differed substantially from that of intestinal epithelial cells. Compared with Caco-2 cells, the number of differentially expressed genes was considerably smaller, suggesting either lower sensitivity or activation of more selective adaptive mechanisms. Nevertheless, alterations involving genes associated with autophagy, oxidative stress and cytoskeletal organization indicate that pancreatic epithelial cells are also capable of responding to bacterial gaseous metabolites. Particularly interesting is the altered expression of *VAMP7*, a regulator of vesicular trafficking and autophagy, together with induction of *SESN2* and *HMOX2*, which are classical markers of oxidative stress adaptation. These findings support the hypothesis that bacterial gaseous metabolites may influence cellular homeostasis independently of direct bacterial invasion (Aoyagi et al. 2018; Russell and Guan 2022; Xin et al. 2026).

Our findings indicate that bacterial gases are biologically active metabolites, not by-products of their metabolism, as until now most researchers have considered H₂ and CO₂ to be end products of fermentation. In this publication, we demonstrate that: bile acids alter the production profile of bacterial gas metabolites; moreover, the resulting gases alter gene expression in Caco-2 and PANC-1 cells. Therefore, the impact of these gases on the human body will depend on their location in the gastrointestinal lumen. This means that gases are becoming another class of microbiome metabolites, similar to short-chain fatty acids (SCFAs). Taken together, these observations suggest that bacterial gaseous metabolites should not be regarded merely as by-products of microbial fermentation but rather as biologically active metabolites capable of influencing host cellular signaling.

Our study has several limitations. First, the experiments were performed using isolates obtained from a single patient, which limits generalization of the observations. Second, only selected bile acids were examined, and additional intestinal metabolites may further influence bacterial gaseous metabolism. Third, although transcriptomic alterations were substantial, the specific gaseous molecules responsible for the observed biological effects remain to be identified. Future studies should include larger collections of clinical isolates, comprehensive metabolomic profiling, and validation in animal models of acute pancreatitis.

Despite these limitations, our work identifies bacterial gaseous metabolism as a previously underexplored component of host–microbiota interactions. The results indicate that gas production depends on bacterial strain and bile acid composition, and that bacterial gaseous metabolites are capable of inducing biologically relevant transcriptional responses in both intestinal and pancreatic epithelial cells. These findings open a new area of research into the role of microbial volatile metabolites in pancreatic diseases and suggest that bacterial gas production may represent a novel mechanistic link between intestinal dysbiosis, altered bile acid metabolism and the pathogenesis of acute pancreatitis. An important future direction will be the identification of patients exhibiting a "gas-producing microbiome phenotype". Such individuals may represent a subgroup at increased risk of altered epithelial signaling and microbiome-associated gastrointestinal or pancreatic diseases. Future studies should determine whether bacterial gaseous metabolite profiles could serve as biomarkers of disease risk or therapeutic response.

In summary, bacterial gaseous metabolites are capable of reprogramming the human transcriptome in a manner comparable to other major microbiome-derived mediators, including lipopolysaccharides (LPS), bacterial toxins, and short-chain fatty acids (SCFAs). Our findings support the concept that bacterial gases constitute a previously underappreciated class of signaling molecules involved in host–microbiome communication.

## Author Contributions

MS: Conceptualization, Funding acquisition, Project administration, Supervision, Writing– original draft, TK: Investigation, Methodology, Validation, Writing – original draft, KM: Methodology, Writing – review & editing, AS: nvestigation, Writing – review & editing, EG: Data curation, Investigation, Methodology, Visualization, Writing – original draft, Writing – review & editing.

## Conflicts of Interest

The authors declare no conflict of interest.

## Acknowledgements

Not applicable.

## Funding

This research was funded by the National Science Center Poland, under Grant No:2018/31/B/NZ6/02472 and by Jagiellonian University Medical College under Grant no: N41/DBS/000699 and N41/DBS/00459.

## Data Availability

All data generated or analyzed during this study are included in this published article.

## Declarations

### Ethics approval and consent to participate

The study was conducted in accordance with the Declaration of Helsinki and approved by the Bioethics Committee of the Jagiellonian University (opinion number KBET 1072.6120.279). Written informed consent for all participants was obtained.

### Consent to publish

Not applicable.

### Competing interests

The authors declare no competing interests.

### Data availability statement

The raw data supporting the conclusions of this article will be made available by the authors, without undue reservation.

### Declaration of Generative AI and AI-assisted technologies

During the preparation of this work, the author(s) used Chat GPT for English language proofreading and grammar correction . The author(s) reviewed and edited the output as needed and take full responsibility for the content of the published article.

